# Denitrifier activity drives diurnal N_2_O flux dynamics in a thawing arctic permafrost

**DOI:** 10.1101/2025.11.04.686510

**Authors:** Dhiraj Paul, Wasi Hashmi, Nathalie Ylenia Triches, Matej Znaminko, Henri M.P. Siljanen, Ivan Mammarella, Sara Hallin, Mathias Göckede, Christina Biasi, Maija E Marushchak

**Affiliations:** Department of Environmental and Biological Sciences, University of Eastern Finland, Kuopio Campus, P.O. Box 1627, FI-70211 Kuopio, Finland; Max Planck Institute for Biogeochemistry, Jena, Germany; Institute for Atmospheric and Earth System Research (INAR)/Forest Science, Faculty of Agriculture and Forestry, University of Helsinki, Helsinki, Finland; Institute for Atmospheric and Earth System Research (INAR)/Physics, Faculty of Science, University of Helsinki, Helsinki, Finland; Department of Forest Mycology and Plant Pathology, Swedish University of Agricultural Sciences, Uppsala, Sweden; Science for Life Laboratory (SciLifeLab), Swedish University of Agricultural Sciences, Uppsala, Sweden; Institute of Ecology, University of Innsbruck, Innsbruck, Austria

**Keywords:** Denitrification, diurnal N_2_O fluxes, metagenomics, metatranscriptomics, nitrous oxide, N_2_O consumption, N_2_O emission, palsa mire, permafrost

## Abstract

Diurnal variation of nitrous oxide (N_2_O) fluxes was recently shown to occur across terrestrial ecosystems, typically with higher daytime emissions. However, the underlying microbial mechanisms remain poorly understood. Here, we investigated the microbial drivers of daily N_2_O dynamics in a permafrost palsa mire, where a higher net N_2_O sink has been observed during dark than under light conditions. By integrating day–night flux measurements with molecular analyses, we could directly link short-term temporal patterns of N□O exchange with diurnal shifts in microbial activity. Quantitative PCR revealed the presence of denitrification genes *nirK* and *nirS*, associated with N□O emissions in soils, and *nosZ* clades I and II, involved in N_2_O consumption. However, only *nirK* and *nosZ* clade I were transcriptionally active. The diurnal pattern of *nirK*/*nosZ* gene expression largely aligned with the diurnal pattern of N_2_O fluxes, with elevated ratios during daytime, particularly in wet and productive fen habitats. Metatranscriptomic analyses further revealed higher daytime expression ratios of N□O-producing relative to N□O-reducing genes ((p450*nor* + *norB* + *hao*)/*nosZ*), in addition to high *nirK*/*nosZ* expression ratios. Members of Alphaproteobacteria, particularly *Rhizobiales* and *Bradyrhizobium*, consistently expressed both *nirK* and *nosZ* across the diel cycle. Analysis of 1,753 metagenome-assembled genomes obtained from the present and previous work further indicate that partial denitrifiers are key players in regulating N_2_O fluxes in arctic permafrost palsa mire. Together, our results provide new insights into the microbial basis for diurnal N_2_O dynamics, which should be considered when estimating ecosystem-climate feedback in terrestrial ecosystems in the Arctic and beyond.

## Introduction

Nitrous oxide (N□O) is a significant greenhouse gas (GHG), with a global warming potential nearly 300 times greater than that of carbon dioxide (CO ) over a 100-year period and the main contributor to ozone depletion [1]. Microbial nitrogen (N) transformation processes in soils represent the dominant source of N□O, accounting for roughly two-thirds of the global emissions [2]. About half of these originate from managed soils influenced by fertilizer inputs and half from natural terrestrial ecosystems. Among the unmanaged soils, tropical soils show the highest emissions due to their rapid N turnover rates, followed by temperate forest soils [3]. Recent work also demonstrates substantial N□O emissions across northern permafrost regions [4]. Despite these advances, N□O fluxes exhibit high temporal variability, which introduces considerable uncertainty in the global N□O emission estimates [5–7]. Most N_2_O flux measurements are so far derived from the summer season, typically using opaque chambers during daytime and at sub-daily sampling frequencies. As a result, diurnal variability and the underling biotic and abiotic mechanism of N_2_O emissions have largely been overlooked. This is a major limitation, as diurnal variation in N_2_O flux appears widespread across ecosystems according to a recent meta-analysis of autochamber data from natural and managed soils [7]. They found that around 80% of studies reported significant diurnal variability, often with higher daytime then nighttime emissions. However, despite microorganisms being central to both N_2_O production and consumption [8], no studies have examined the microbial mechanisms driving patterns of diurnal fluctuations.

Our recent study of N□O fluxes at Stordalen Mire, a arctic permafrost ecosystem, shows diurnal patterns with a higher net N_2_O sink during dark periods [9], suggesting higher N_2_O consumption relative to N_2_O production at night. The only known pathway for N_2_O uptake or consumption is through microbial-mediated reduction of N_2_O to N_2_. This process is catalyzed by the N_2_O reductase, encoded by the *nosZ* gene and phylogenetically divided into two distinct clades: Clade I and Clade II [10, 11]. Clade II *nosZ* has been shown to be particularly important in controlling soil N_2_O emissions [12, 13]. By contrast, denitrification, the step wise reduction of nitrate to N□O or dinitrogen gas (N_2_), is the dominant source of N□O in soils [14, 15]. The key reaction in denitrification leading to gaseous N species formation is the reduction of nitrite to nitric oxide (NO), which is catalyzed by nitrite reductases encoded by *nirK* and *nirS* genes [16]. Nitric oxide then can be further reduced to N_2_O by a number of different NO reductases. Notably, denitrification is largely a community trait with different reactions performed by community members specialized in one or several reactions in the denitrification pathway [17, 18].

Based on the current understanding of the microbial processes at play, we hypothesize that the higher N_2_O uptake in the night and the higher net N□O emission during the day would be explained by the relative differences in expression of *nosZ* genes with those in preceding steps the denitrification pathway. To test our hypothesis and determine microbial drivers of diurnal N□O dynamics, we measured field N_2_O fluxes at Stordalen Mire in four habitats along a thawing permafrost gradient during day and at night with transparent chambers, preserving the natural light conditions, and combined these measurements with microbial sampling and determination of underlying biotic and abiotic factors. We expected the highest diurnal variation in habitats with higher content of reactive N and DOC supporting the denitrifier communities. The microbial functional analyses with a focus on denitrifier gene expression and analyses of metagenome assembled genomes (MAGs) were based on a combination of quantitative PCR, metatranscriptomics, and metagenomics. This integrative approach provides novel mechanistic insights into diel microbial regulation of net N□O emissions and underscores the importance of resolving diurnal processes to better estimate microbial contributions to permafrost region and global scale N□O budgets.

## Materials and methods

### Study site and sample collection

Our study was conducted at Stordalen Mire (68°35’N, 19°04’E), a permafrost peatland located 10 kilometers east of the Abisko Natural Sciences Research Station (Fig. 1A-E). The region lays in the southern border of permafrost occurrence, and isolated permafrost is currently present in the dry uplifted areas on the peatland (palsa), but it is threatened by ongoing permafrost thaw [19]. A detailed description of the sampling site is provided in the supplementary (Supplementary methods)

**Figure 1:**
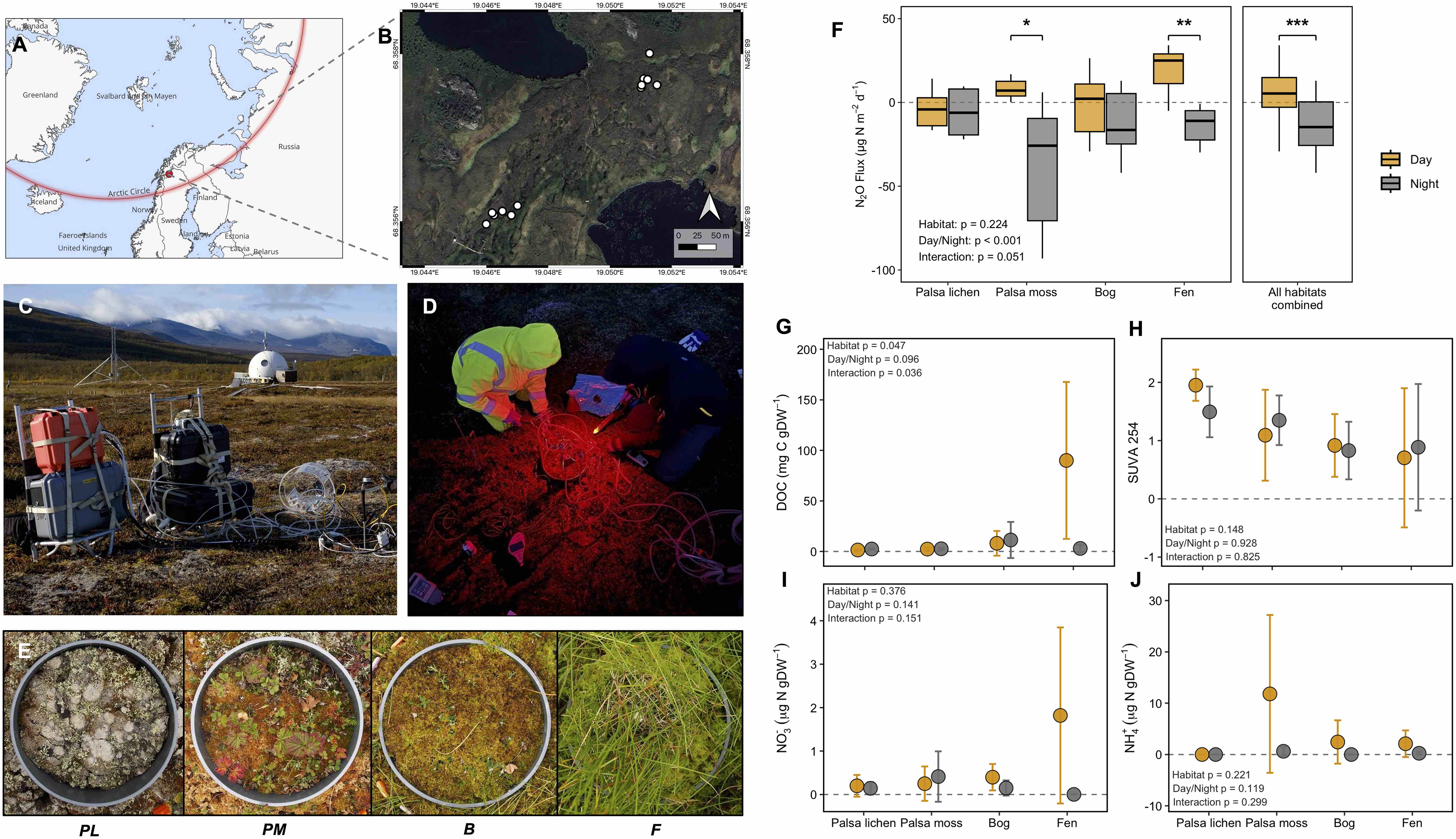
Sampling location, diurnal N_2_O emissions, and soil chemistry in four different habitats in an arctic permafrost peatland. (A,B) Sampling location in Abisko, Sweden, and photos illustrating field sampling conducted during (C) daytime and (D) nighttime, respectively. (E) Four different habitats based on the dominating plant species were selected for N_2_O flux measurements and sampling: Palsa lichen (PL), Palsa moss (PM), Bog (B) and Fen (F), along a permafrost thaw gradient (Photos C-E by W. Hashmi). (F) Boxplots show daytime and nighttime N□O fluxes across the four habitats (n=6 per habitat) and all habitats combined (n=48). Boxes represent lower and upper quartiles, median (thick black line), smallest and largest values without outliers (thin black line); (G-J) Dot-and-whisker plot showing differences in DOC (dissolved organic carbon), SUVA_254_, NH_4_ (ammonium) and NO_3_ (nitrate) at 2 cm soil depth at the different habitats (mean ± SD, *n*=3). The dashed horizontal line marks the null expectation (0), where values above and below the line indicate positive and negative deviations, respectively. Statistical differences in fluxes and soil chemistry among habitats were assessed using two-way ANOVA, and paired t-test was applied to assess day-night differences within each habitat. Asterisks indicate significant differences between day and night (*t*-test, ∗*P* <0.05, ∗∗*P* <0.01, ∗∗∗*P* <0.001). Gold and gray colors indicate daytime and nighttime sampling, respectively.

We selected a subset of 12 chamber flux plots in two transects on a dry to wet thawing gradient, with three replicates for each of four habitat types: palsa lichen, palsa moss, bog, and fen. These flux plots represent a subset of the 24 plots studied by Triches et al. [9] and were selected based on representativeness of the spatial variability within the mire and accessibility during night-time measurements. During a field campaign at Stordalen Mire in mid of August, 2024, we conducted *in situ* N_2_O flux measurements in comparable conditions in the day and the following night (Fig. 1C-D) and collected surface peat samples for microbiological and geochemical analysis. We chose this time during the late growing season because of the reoccurrence of dark nights after the polar day. Soil samples were collected from both 2 cm and 15 cm depths. For microbiological sampling, samples were collected in triplicate from the vicinity of each flux plot using a sterile forceps and pooled into a single sample per depth in a sterile falcon tube pre-treated with DEPC (Diethyl pyrocarbonate)-water. A total 48 samples were collected, comprising two sampling times (diurnal sampling), four habitats, two depths, and three biological replicates. Samples were rapidly flash-frozen in liquid N_2_ in the field, and transported on dry ice to our laboratories, where they were stored at −80 °C. Simultaneously, we took samples for geochemical analysis which were stored in sterile falcon tubes, placed in a cooler for transport and stored at +4 °C until analysis.

### N_2_O flux measurements and auxiliary data

N_2_O fluxes for palsa lichen, palsa moss, bog, and fen were measured with the manual chamber method previously used for extensive flux measurement at Stordalen Mire over three growing seasons 2022-2024 and documented in detail by Triches et al. [9]. Our manual chamber fluxes were recorded during daytime (11:00–18:00) and nighttime (22:00–03:00) using a transparent chamber and portable gas analyzers MIRA Ultra N_2_O/CO_2_ (Aeris, CA, USA) by default, with Li-7820 (Licor, NE, USA) as a backup instrument. In addition to N_2_O fluxes, we recorded also CO_2_ and CH_4_ fluxes using a Li-7810 (Licor; NE, USA) analyzer. Chamber temperature, air pressure, humidity, PAR (photosynthetically active radiation), soil temperature, and moisture were also recorded. Flux measurement details are given in the supplementary (Supplementary methods). Throughout the article, we use the sign convention where net emission from the ecosystem is reported as positive fluxes and net atmospheric consumption as negative fluxes.

### Soil chemical properties

Soil characteristics were determined within one week from the sampling from peat samples taken during both day and night flux measurement (n=3). The samples were homogenized, and large roots and above ground vascular plants were removed. Soil pH and electrical conductivity (EC) were measured in a soil slurry (a soil-to-water ratio of 3:5 (v/v)). Water extractable content of ammonium (NH_4_^+^), nitrate (NO_3_^-^), dissolved organic carbon (DOC) and dissolved nitrogen (DN) were analyzed from water extracts using a soil-to-water ratio of 3:10 (v/v), and dissolved organic nitrogen (DON) was calculated as a difference of DON and mineral N. Total C, N, C:N ratio, δ^13^C, and δ^15^N were measured from dried soil samples after homogenization by a ball mill. Photometric measurements of SUVA_254_ were used to estimate DOC aromaticity. The chemical analyses are described in detail in the supplementary (Supplementary methods).

### DNA and RNA extraction

Total community DNA and RNA were extracted from soil samples using the RNeasy PowerSoil RNA Extraction Kit (Qiagen, Germany), followed by the Soil DNA Elution Kit (Qiagen, Germany), according to the manufacturer’s protocol with minor modification. In brief, to separate out the phenol and aqueous phase properly, 1.5 g of soil was used instead of the 2 g recommended in the kit protocol. This modified protocol was based on our test extractions and resulted in a better DNA and RNA yield from the sample material used in this study. The quality of the RNA and DNA was assessed using a NanoDrop spectrophotometer and the quantity was determined using Qubit High Sensitivity (HS) RNA and HS DNA Assay Kits (Thermo Fisher Scientific, MA, USA), respectively.

For quantitative PCR (qPCR) of expressed genes, a portion of the RNA was treated with DNase treatment followed by complementary DNA (cDNA) synthesis. Removal of residual DNA from the eluted RNA was confirmed by measuring DNA concentrations using a Qubit fluorometer after DNase treatment and by performing PCR amplification of the 16S rRNA gene using universal primers 341F and 518R [20] prior to cDNA synthesis. After confirming that there was no amplification, we converted the RNA into single-stranded cDNA. Both DNA and cDNA samples were stored at −20 °C and RNA samples were stored at −80 °C.

### Abundance and expression of denitrification genes determined by qPCR

Quantitative real-time PCR (qPCR) was used to quantify key functional genes involved in denitrification (NO-forming nitrite reductase genes *nirS* and *nirK*) and N_2_O consumption (*nosZ* clade I and clade II), along with bacterial and archaeal 16S rRNA genes to assess the total bacterial/archaeal abundance. Detailed PCR conditions are provided in Supplementary Table S1. To determine the total copy number and expression of N-cycling genes, we performed qPCR using both DNA and cDNA as templates, respectively. Each qPCR reaction was conducted with a total volume of 10 µL, including three biological replicates, using the GOTAG SYBR Green qPCR kit on Bio-Rad MyCycler system (CA, USA). Each reaction contained 5 ng of template DNA and 0.5–1.0 µM of each primer to ensure accuracy and consistency. To test for potential PCR inhibition, we amplified a known amount of the pGEM-T plasmid (Promega, WI, USA) with plasmid-specific T7 and SP6 primers added to the 5 ng of extracted DNA, as well as non-template controls. No inhibition of the PCR reaction was detected based on comparison of cycle threshold (Ct) values between DNA extracts and non-template controls for the amount of template used. The amplifications were verified by melting curve analyzes and agarose gel electrophoresis.

### Metatranscriptome sequencing and assembly

For metatranscriptome analysis, extracted community RNA was treated twice with TURBO DNase (Invitrogen) to remove all DNA. Following DNase treatment, we checked absence of DNA with Qubit readings, further confirmed by absence of DNA amplification by 16S rRNA gene [20]. Ribo-zero rRNA Removal Kit (Epicentre, WI, USA) was used to remove ribosomal RNA (rRNA) from the total RNA extract. A cDNA library was constructed and subsequently sequenced using the 2 x150 bp chemistry on NovaSeq X Plus Series (150-bp paired-end) at Novogene, Germany. Raw reads were quality-checked with FastQC [21]. High-quality reads (Q > 30) were trimmed (Trimmomatic) [22] and rRNA removed (SortMeRNA) [23]. About 1.35 billion non-rRNA reads were assembled (rnaSPAdes) [24], ORFs predicted (Prodigal) [25], and functionally annotated (eggNOG-mapper v2.1.12) against KEGG, COG and pfam database. Reads were mapped (Bowtie2) [26], alignments processed (SAMtools) [27] and expression levels normalized as Transcripts Per Million (TPM). Taxonomy assignment of expressed *nosZ* and *nir* gene were determined with GraftM [28] using existing phylogenies [29, 30]. A detailed description of the analysis is available in the supplementary (Supplementary methods).

### Metagenome sequencing, assembly, binning and functional annotation

From the present study, extracted community DNA was used for DNA fragmentation and library construction using the NEBNext Ultra II DNA Library kit (New England Biolabs, MA, USA). The quality and quantity of the fragment library were assessed using the Qubit dsDNA HS assay kit (Thermo Fisher Scientific, MA, USA) and the Agilent 2200 TapeStation (Agilent, CA, USA), respectively. A high-quality library was normalized, pooled, and subsequently sequenced using 2×150 bp chemistry on the NovaSeq X Plus Series (Illumina) at Novogene, Germany. Raw reads were quality-checked (FastQC), trimmed (Trimmomatic), and assembled (MEGAHIT) [31]. Genomes were binned (MaxBin2 [32], MetaBAT2 [33], CONCOCT [34]), consolidated and dereplicated (DASTool, dRep) [35, 36], quality-checked (CheckM) [37], and taxonomically classified (GTDB-Tk) [38]. In total, 282 MAGs were recovered. Detailed description of the analysis is provided in the supplementary. (Supplementary methods).

To enhance our understanding of the microbial control on the N-cycle in the mire ecosystem, we undertook a comprehensive MAGs analysis by integrating the 125 MAGs recovered in this study (having 70% completeness and less than 10% contamination) [37], with an existing Stordalen Mire MAG database comprising 13,290 MAGs generated through the EMERGE project [39, 40]. The database containing MAGs with same quality criteria (70% completeness and less than 10% contamination). More information about the constructed database is available in McGivern et al. (2024) [40]. For this study, we were only interested in the 1,628 MAGs reconstructed from metagenomic data collected from depths of 0 to 15 cm in the palsa, bog, and fen habitats to match with the current data set. As a result, around 1,753 MAGs, combining data from our study and previous studies, were considered for functional analysis.

To identify functional genes associated with nitrogen fixation, nitrification and denitrification, specifically *nifH, amoA, hzoA, narG, napA, nrfA, nxrB, nirK, nirS, norB,* and *nosZ,* we utilized targeted gene hmmer profiles when analyzing the MAGs [41]. This enabled us to search for sequences of each functional gene with a maximum *E*-value cutoff of *E* <0.000001. Gene function and relatedness were confirmed by comparing sequences against UniProt-TrEMBL, Swiss-Prot, and NR databases using the *tblastx* method in the DIAMOND sequence search accelerator. ORFs were predicted from metagenome assemblies using Prodigal (v2.6.3, meta mode) and functionally annotated were carried out using eggNOG, KEGG database. Relative abundance (count per million, CPM) of genes was calculated by mapping reads from each sample with Bowtie2 (v2.2.0) [26]. Detailed description of the analysis is provided in the supplementary (Supplementary methods).

### Statistical analysis

Data are presented as mean ± standard deviation (SD). In situ N□O fluxes were determined from six independent observations (n=6), whereas geochemical and qPCR measurements were obtained from three biological replicates (n=3). For variables subjected to parametric analyses, data distributions for normality were checked using Shapiro–Wilk test, and log-transformation was applied when necessary to meet assumptions of normality and homogeneity of variance. The effects of time of day (day vs. night), habitat type, and their interaction on N_2_O fluxes and soil geochemical variables were evaluated using two-way analysis of variance (ANOVA). Differences between daytime and nighttime measurement, both across all habitats and within individual habitats, were further assessed using *t*-test. Differences in qPCR and omics datasets between day and night, both across and within habitats, were analyzed using *t*-test on log-transformed data. Associations between microbial and geochemical variables were determined using Spearman’s rank correlation analysis. Statistical significance was accepted at *p* <0.05. R packages, such as ggplot2 (v.3.5.2) [42], Reshape2 (v.1.4.4) [43], ggpubr (v.0.6.1) [44], corrplot (v.0.95) [45], tidyr (v.1.2.0) [46], dplyr (v.1.0.9) [47], readxl (v.1.4.0) [48], edgeR (v.3.16) [49], cowplot (v.1.1.1) [50] were used for data analysis and visualizations. All analyses and visualizations were done using R version 4.2.2 (R Core Team, 2024).

## Results

### Diurnal variability in *in situ* N_2_O fluxes

Fluxes of N_2_O varied between sources and sinks, with a significantly higher mean N_2_O flux across all habitats during the day than at night (5.54 ± 16.59 μg N_2_O-N m^−2^ d^−1^ vs. −21.2 ± 29.8 μg N_2_O-N m^−2^ d^−1^; ANOVA, *p*<0.001; Fig. 1F, Supplementary Table S2). The largest, statistically significant differences between day and night fluxes were found in fen (pairwise *t*-test, *p* <0.01) followed by palsa moss (pairwise *t*-test, *p* <0.05), with less diurnal variability in bog, and similar fluxes in day and night in palsa lichen (Fig. 1F). Notably, the fen, representing the wettest habitat and the final stage of the permafrost-thaw gradient from the intact palsa to thermokarst fen, exhibited the highest daytime N□O emissions (19.34 ± 15.14 μg N□O-N m^2^ d^1^) and shifted to net N□O uptake at night (−8.7 ± 15.9 μg N□O-N m^2^ d^1^) (Fig. 1F; Supplementary Table S2).

Consistent with the observed diurnal N□O dynamics, auxiliary measurements revealed habitat effects on CH and CO fluxes (Supplementary Fig. S1; two-way ANOVA, *p* = 0.001 and *p* = 0.096, respectively). Along the thawing gradient the CH_4_ flux increased from net consumption or small emissions in the palsa sites through moderate emissions in the bog to large CH_4_ release in the fen. Similarly, we observed an increase in nighttime CO_2_ emission and daytime net CO_2_ sink along this gradient, resulting in the highest diurnal amplitude in CO_2_ fluxes in the fen. Overall, diurnal variability was observed in both CO_2_ fluxes and CH_4_ fluxes (Supplementary Fig. S1). PAR values were significantly (*t*-test; *p* = 0.003) higher during the daytime measurements than at nighttime (561 ± 141 μmol m^−2^ s^−1^ and 0.1 ± 0.14 μmol m^−2^ s^−1^, respectively; Supplementary Table S2). No significant differences (*t*-test, *p*>0.05) between day and night measurements were observed in soil temperature at 15 cm in any of the habitats (Supplementary Table S2). The mean soil temperature at 15 cm across all habitats was 7.8 ± 1.74 and 8.4 ± 1.1 °C in the day and at night, respectively. Similarly, volumetric soil moisture content did not differ significantly between day and night (*t*-test, *p*>0.05), with values of 50.7 ± 6.4% during the day and 50.2 ± 9.4% at night (Supplementary Table S2).

### Geochemical characterization of peat

Geochemical characteristics of surface peat were overall similar among habitats, with consistently low pH (∼4) and low electrical conductivity (EC) ranging from 10 to 50 µS cm^−1^ (Supplementary Table S3). C:N ratios were frequently elevated (>25–100) across the habitats, indicating more carbon-rich and nitrogen-poor dissolved organic matter (Supplementary Table S3), suggesting relatively low N availability in the system. A few, notable differences were observed in water soluble C and N pools among the habitats (Fig. 1G-J, Supplementary Figs. S2-S4, Supplementary Table S3). Dissolved organic nitrogen (DON) was higher in fen, particularly 15 cm depth, but remained lower in palsa and bog habitats at both the depth, indicating limited soluble organic N pools (Supplementary Fig. S4 and Fig. 1). Fen also had significantly higher concentration of DOC (ANOVA*, p* = 0.047 at 2 cm and *p* = 0.041 at 15 cm) and lower DOC aromaticity indicated by a lower SUVA compared with other mire habitats (Fig. 1 and Supplementary Fig. S2). The water-extractable NH_4_^+^ at 2 cm was higher in palsa lichen than in other habitats (Fig. 1J). Although day–night differences in DOC were not statistically significant (ANOVA*, p*>0.05), marginal diurnal variation was noted at 2 cm depth, as well as for both DOC and DON at 15 cm depth in fen habitats, with higher concentrations noted during the daytime than at night (Fig. 1 and Supplementary Fig. S2 and S4). Correlation analyses revealed predominantly positive but non-significant associations between N□O flux and PAR, DOC, DON, C/N ratio, and NO at 2 cm depth (*p*>0.05; Supplementary Fig. S5A). At 15 cm depth, N□O flux was also positively but non-significantly related to NO and C/N (*p*>0.05) but showed a significant positive association with DOC (ρ = 0.73, *p* = 0.01) and DON (ρ = 0.75, *p* = 0.007), likely driven by the influence of fen habitats (Supplementary Fig. S5B). The lack of a significant PAR and N□O relationship, despite similar diurnal patterns, suggests that short-term temporal variability is governed by different processes than those controlling spatial variation across habitats.

### Diurnal pattern of N-cycling gene abundance and expression

Quantitative PCR analyses showed that targeted genes involved in N□O reduction (*nosZ* clades I and II) and nitrite reduction in denitrification (*nirK* and *nirS*) were present at the DNA level in all habitats at both soil depths. However, no significant differences (*t*-test, *p* > 0.05) were observed between daytime and nighttime sampling for the abundance total microbial communities (Supplementary Fig. S6) or abundance of denitrification genes (Supplementary Fig. S7). The abundance of *nirK* was higher than that of both *nirS* and *nosZ* clades I and II at both depths and *nirS* gene was mainly prevalent in the fen compared with the other habitats (Supplementary Fig. S7).

At the RNA level, there were differences in denitrifier gene expression between day and night. The gene expression ratio of *nirK* to *nosZ,* a proxy for gaseous N loss in the form of N_2_O *vs*. N_2_ during denitrification, aligned with the flux patterns. Across all habitats, the *nirK/nosZ* expression ratio was 5-20-fold higher during the day than at night (*p* <0.05; Fig. 2A, B) at 2 cm soil depth but not at 15 cm (Supplementary Fig. S8A, B). The diurnal difference was significant in the fen and palsa moss habitats (*t*-test, p < 0.05; Fig. 2A), consistent with the habitat-specific patterns observed for N□O fluxes (Fig. 1F). The relationship between N□O flux and the *nirK*/*nosZ* ratio at 2 cm across all habitats was not significant (Spearman’s ρ = 0.02, *p* = 0.92; Supplementary Fig. S5A), suggesting that although the expression ratio was an important predictor for diurnal N_2_O variability within individual habitats, it poorly explained habitat differences in the diurnal amplitude.

**Figure 2:**
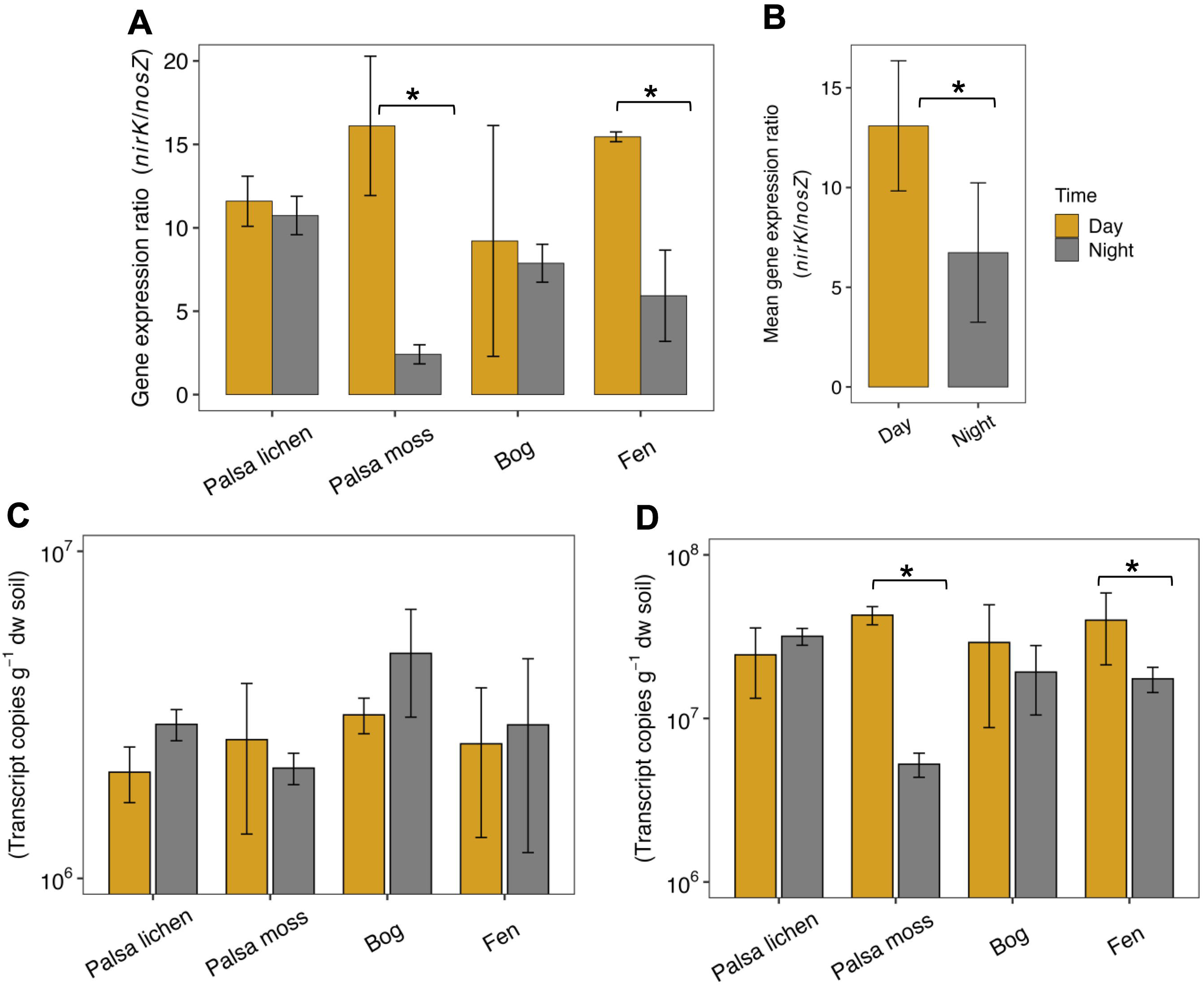
Expression of N□O reductase and NO-forming nitrite reductase genes in the four habitats (Palsa lichen, Palsa moss, Bog and Fen) based on qPCR analysis of soils at 2 cm depth. (A) Habitat-specific and (B) overall *nirK*/*nosZ* expression ratios, representing the balance between the potential for N□O production and N□O reduction. (C) Expression of N□O reductase gene *nosZ* clade I and (D) expression of nitrite reductase gene *nirK*. Three biological replicates were analyzed for each habitat, with each biological replicate representing the mean of two technical replicates. Data are presented as mean ± standard deviation (SD). Gold and gray indicate daytime and nighttime samples, respectively. Differences in gene expression and *nirK/nosZ* expression ratios between daytime and nighttime samples, both within individual habitats and across all habitats were assessed using *t*-tests. Asterisks denote significant differences between day and night expression (∗*P* < 0.05).

For genes encoding nitrous oxide reductase, we only observed expression of *nosZ* clade I but not *nosZ* clade II despite its presence at DNA level. In accordance with the stronger net N_2_O sink during nighttime, *nosZ* clade I expression at 2 cm tended to be higher at night than in the day, although not significantly higher (*t*-test, *p* > 0.05, Fig. 2C) and the opposite trend was observed at 15 cm depth (Supplementary Fig. S8C). For genes coding for nitrite reduction in denitrification, we only observed expression of *nirK* across all habitats and soil depths (Fig. 2D, Supplementary Fig. S8D). Despite testing various template concentrations, *nirS* expression was not detected, even though it was present at the DNA level (Supplementary Fig. S7). The diurnal pattern of *nirK* expression showed significantly higher gene expression at 2 cm soil depth during daytime in the fen and palsa moss (Fig. 2D, *t*-test, *p* <0.05) but overall, *nirK* expression showed non-significant (*p* >0.05) positive associations with N□O fluxes (Supplementary Fig. S5).

### Metatranscriptomic and metagenomic analyses

The metatranscriptome analysis confirmed the overall diurnal pattern of *nirK*/*nosZ* expression ratio obtained by qPCR (Fig. 3A,B), suggesting that relative differences between denitrification genes at the transcriptional level control day and night N_2_O flux fluctuations in N_2_O. This is further supported by the expression ratio of N_2_O production *vs*. N_2_O reduction genes ((p450*nor*+*norB*+*hao*)/ *nosZ*)), which was higher during daytime (Fig. 3C,D). The actively transcribing *nirK* community was dominated by bacteria, with members of the Alphaproteobacteria, particularly *Rhizobiales* and *Bradyrhizobium,* showing high *nirK* expression (Fig. 3E). The composition of the *nirK* expressing community showed minimal variation between day and night (Fig. 3E). Similarly, *nosZ* was expressed by bacteria, with notable contributions from *Bradyrhizobium* and diverse members of *Rhizobiales*, following similar taxonomic patterns as those of *nirK* expression (Fig. 3E). These taxa were also abundant at the metagenome level (Supplementary Table S4). Similar to the qPCR results, our metatranscriptomics analysis also did not detected expression of *nirS* (cytochrome cd1 nitrite reductase) and its associated genes (*nirF*, *nirN*, *nirJ*, *nirE*, *nirT*) (Supplementary Table S5).

**Figure 3:**
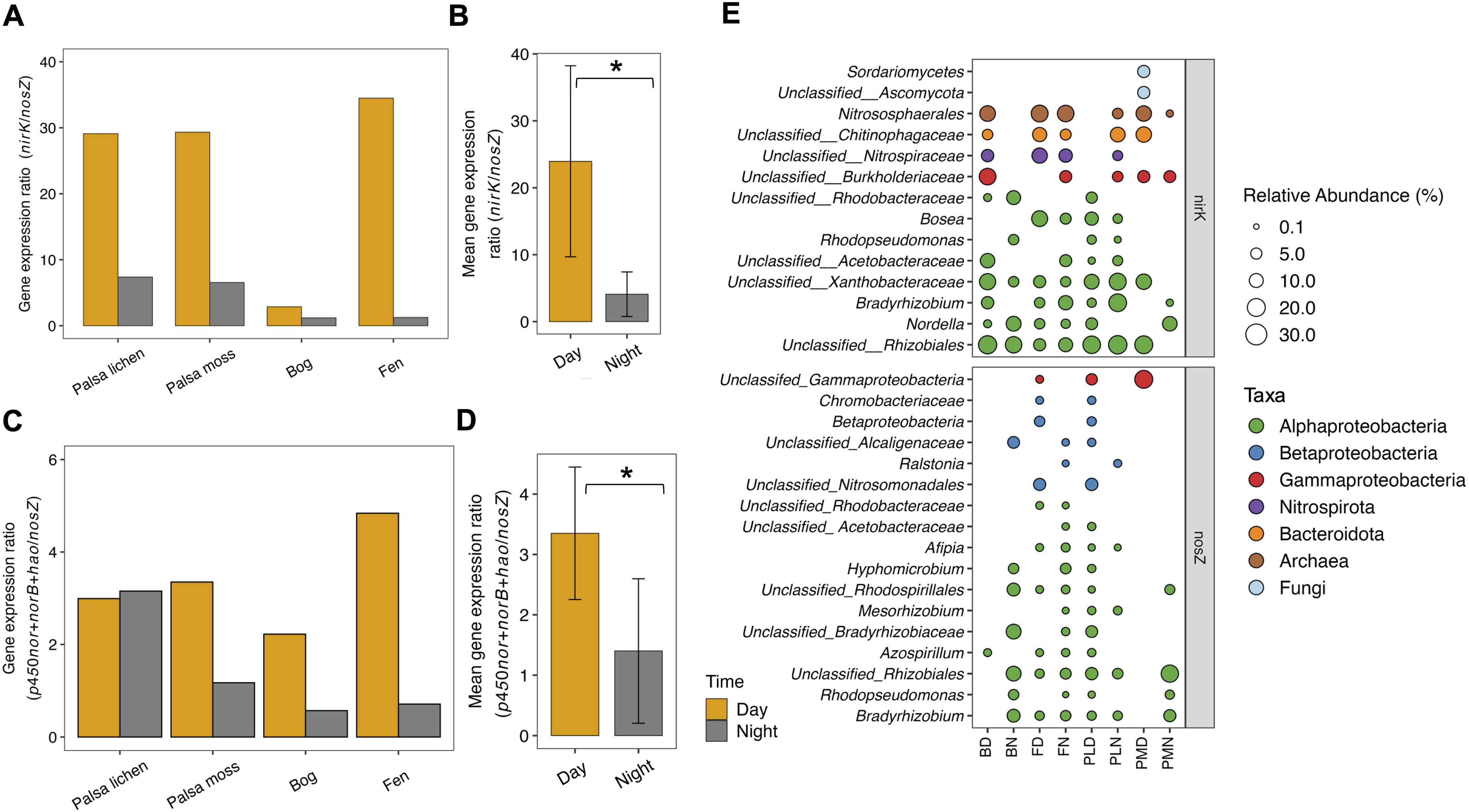
Day–night ratios of expressed N□O producing and reducing genes and their taxonomic distribution based on metatranscriptomic analysis of soils at 2 cm depth. (A) Denitrifier nitrite reduction-to-N□O consumption gene ratio (*nirK*/*nosZ*) across the four habitats (Palsa lichen, Palsa moss, Bog, and Fen) and (B) overall *nirK/nosZ* ratio across all habitats combined (mean ± SD, *n*=4). (C) N□O production-to-consumption gene ratio ((p450*nor*+*norB* +*hao*)/*nosZ*) across habitats and (D) overall ((p450*nor*+*norB* +*hao*)/*nosZ*) across all habitats combined (mean ± SD, *n*=4). Statistical significance between daytime and nighttime gene expression ratios across habitats were assessed using a *t*-test. Asterisks denotes significant differences between day and night expression (∗*P* < 0.05). Gold and gray indicate daytime and nighttime samples, respectively. (E) Taxonomic distribution of *nirK* and *nosZ* transcripts in diurnal samples from the four habitats. Sequences that could not be classified at the genus level are labeled “unclassified” followed by the family name. The size of each circle indicates the relative abundance of each taxon. Abbreviations: BD: bog daytime; BN: bog nighttime; FD: fen daytime; FN: fen nighttime; PLD: palsa lichen daytime; PLN: palsa lichen nighttime; PMD: palsa moss daytime; PMN: palsa moss nighttime.

Among the 1,753 MAGs, the denitrification genes *nirS*, *nirK*, *norB*, and *nosZ*, were detected in 23, 345, 368, and 88 MAGs, respectively (Fig. 4A, Supplementary Table S6). Among MAGs, we found high (*nirK*+*nirS*)/*nosZ* ratios, indicating a greater genomic potential for N_2_O production than reduction (Fig. 4A,B). Notably, no single MAG contained nitrification genes coding for ammonia oxidation (*amoA*) (Fig. 4A) and metagenomic read-based analyses support this trend, revealing a limited presence of reads associated with ammonia-oxidizing bacteria and archaea (Fig. 4C, Supplementary Table S7). This was further confirmed by metatranscriptome analysis, which revealed no detectable expression of *amoA* gene and overall low expression of hydroxylamine oxidoreductase (*hao*) (2.7 TPM) (Supplementary Table S5). By contrast, higher expression of *norB* (21.2 TPM) in bacterial and p450*nor* (4.1 TPM) in fungal denitrification pathways was observed (Supplementary Table S5), suggesting N_2_O production from denitrification dominated compared to nitrification in this system.

**Figure 4:**
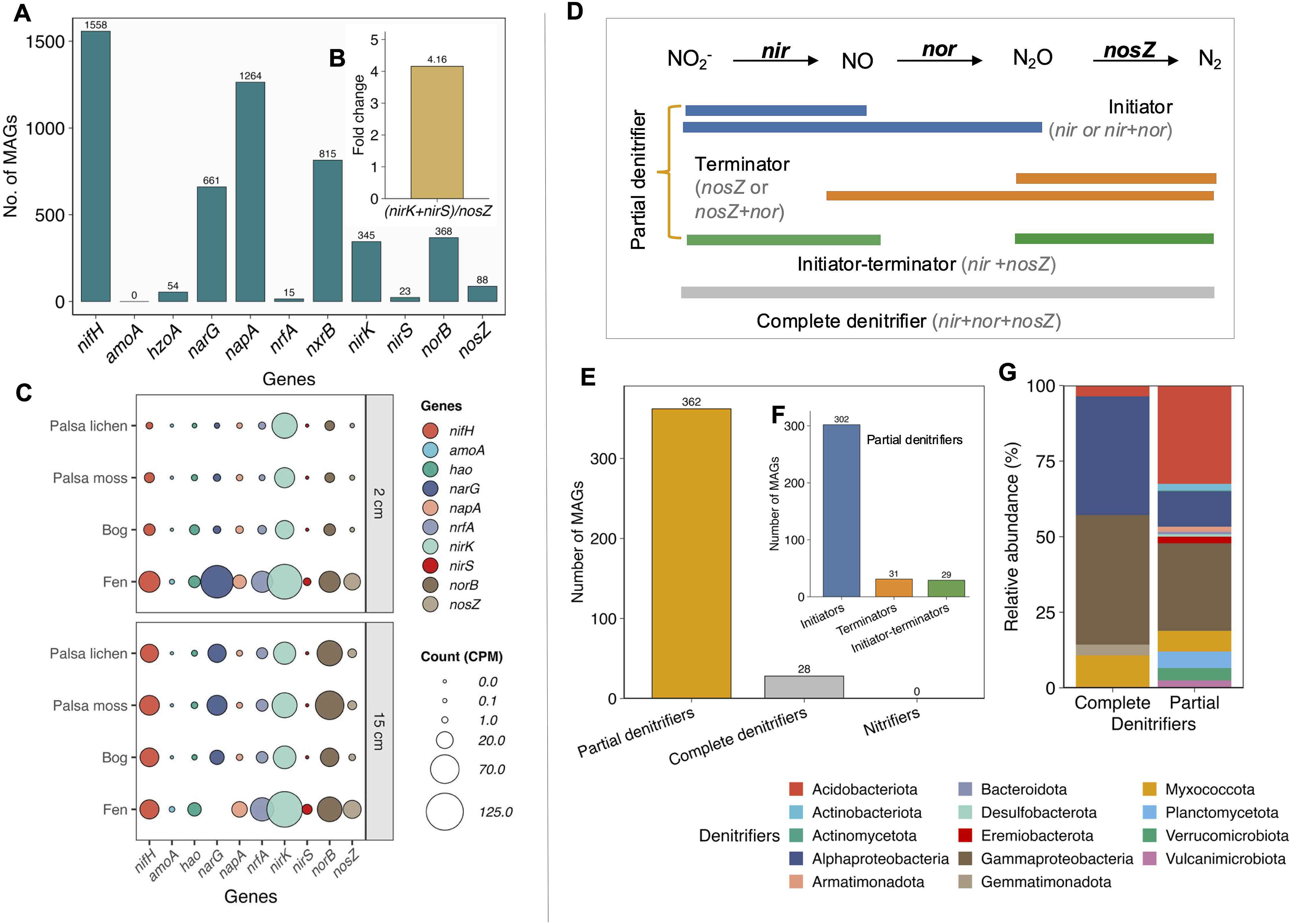
Genome reconstruction from metagenomes and presence of nitrogen cycling genes. (A) Prescence of nitrogen cycling genes across metagenome-assembled genomes (MAGs) obtained in the present study and from Li et al. (2025). (B) Fold change in abundance of *nir* vs *nosZ* genes across the MAGs. (C) Relative abundance of nitrogen cycling genes in the four habitats at 2 and 15 cm depth from the present study. Circle size is proportional to count per-million, and colours denote individual nitrogen cycling genes. (D) Classification of denitrifier types based on Pold et al (2025) with complete denitrifiers and partial denitrifiers divided into initiators (*nir* with or without *nor*), terminators (*nosZ* with or without *nor*) and initiator-terminators (*nir* and *nosZ*). (E) Distribution complete denitrifiers, partial denitrifiers, and nitrifiers across MAGs. (F) Distribution of partial denitrifier types. (G) Taxonomic composition of partial and complete denitrifiers represented by MAGs. Gene abbreviations: *nifH*, nitrogenase reductase; *amoA*, ammonia monooxygenase subunit A; *hzoA*, hydrazine oxidoreductase A; *narG*, nitrate reductase alpha subunit; *nxrB*, nitrite oxidoreductase beta subunit; *nrfA*, cytochrome c nitrite reductase; *napA*, periplasmic nitrate reductase; *nirK*, copper-containing nitrite reductase; *nirS*, cytochrome cd_1_ nitrite reductase; *norB*, nitric oxide reductase; *nosZ*, nitrous oxide reductase.

Based on the framework of Pold et al. (2025) [18], MAGs were further categorized denitrifier types in terms of complete denitrifiers (those with *nir*, *nor*, and *nosZ* genes) and partial denitrifiers (those lacking either *nir* or *nos*Z). Partial denitrifiers further divided into initiators (*nir* with or without *nor*), terminators (*nosZ* with or without *nor*) and initiator-terminators (*nir* and *nosZ*) (Fig. 4D). We identified 28 genomes, representing 7.1% of the MAGs, that were complete denitrifiers from five bacterial phyla (Fig. 4E; Supplementary Table S8), with the most prevalent being Pseudomonadota (dominated by the classes Alphaproteobacteria, Gammaproteobacteria, each accounting for 40%), followed by Myxococcota (10%) and Acidobacteriota (Fig. 4G). By contrast, partial denitrifiers dominated, comprising 365 MAGs (Fig. 4E, F) and belonging primarily to Pseudomonadota (Alpha- and Gammaproteobacteria), Acidobacteriota, and Planctomycetota (Fig. 4G; Supplementary Table S8).

## Discussion

By combining diurnal N□O flux measurements with molecular analyses of N-transformations, especially those by the denitrifier community, we present a mechanistic link between temporal shifts in gene expression and N□O dynamics. Nitrous oxide emissions from Stordalen Mire were consistently lower at night than during the day, corroborating earlier observations from the same site [9], and other terrestrial ecosystems [7]. Along Stordalen Mire thaw gradient, the strongest day– night contrasts occurred in the fully thawed fen, which functioned as the largest daytime N_2_O source and strong nighttime sink. The second highest significant diurnal variability was found in palsa moss, while the diurnal amplitude was smaller in the bog, and negligible in the palsa lichen. This highly dynamic N_2_O source-sink pattern in the fen implies active turnover of mineral N and complements previous understanding of enhanced greenhouse gas turnover as CO_2_ and CH_4_ with thaw-induced inundation [51]. Substantial N_2_O production in the fen, being the final stage of a permafrost thawing gradient, has been previously reported [52], but our observations of increasing N_2_O sink activity during night gives a novel perspective on N_2_O fluxes from these habitats [9].

The diurnal N□O dynamics in the permafrost peatland habitats were associated the *nirK/nosZ* expression ratio, a proxy for gaseous N losses in the form of N_2_O vs N_2_ [13, 53, 54]. This provides molecular evidence that diurnal variability in N□O fluxes in sub-arctic peatlands [9] is driven by microbial activity, particularly by denitrifying microorganisms. Daytime peaks in N□O fluxes corresponded with elevated *nirK/nosZ* ratios driven by increased daytime *nirK* expression, while nighttime minima corresponded to reduced ratios likely resulting from a combination of decreased *nirK* expression and enhanced *nosZ* expression. This diurnal decoupling of N□O production and reduction suggests that enhanced incomplete denitrification during daytime increases the N□O:N product ratio. Consistent with these results, our metatranscriptomic analysis revealed reduced ratios of N□O production to reduction genes ((p450*nor* + *norB* + *hao*)/*nosZ*), which also suggest a shift towards more complete denitrification process at night. Furthermore, higher N□O emission and *nirK* expression during daytime likely reflects increased nitrate and dissolved organic carbon (DOC) availability during daytime, at least in fen sites. Previous studies have also reported positive association between nitrate and DOC availability and *nirK* abundance and expression [55, 56]. Notably, diurnal N□O fluxes matched the *nirK*/*nosZ* expression ratio at the surface (2 cm), but not at 15 cm depth. This suggests limited N□O diffusivity in water-saturated peat and reduction of N□O during upward transport, especially in fens. Consequently, surface-layer (2 cm) gene expression ratios should better predict net atmospheric N□O exchange [57]. This points towards the importance of the living moss layer in driving diurnal variability in N_2_O fluxes, likely via strong light-dependency of photosynthetic activities or influx of N through diazotrophs in the moss layer [9].

Both *nosZ* clades I and II were present in the total gene pool in all four habitats, but only *nosZ* clade I was expressed. *nosZ* clade II is typically associated with phylogenetically diverse non-denitrifying organisms [11], whereas *nosZ* clade I is less diverse, found primarily in Pseudomonadota, and is the most common N_2_O reductase in complete denitrifiers [17, 18]. Consistently, the actively transcribing *nosZ* community was dominated by Pseudomonadota, particularly Alpha- and Betaproteobacteria (Fig. 3E). Previous studies have demonstrated that N□O-reducing communities in nitrate-rich soils (e.g., fertilized croplands) tend to exhibit more *nosZ* clade II than clade I counts when compared with tundra soils, including peatlands [18]. Further, in root associated environments, clade I dominates [58], altogether suggesting habitat-specific niche differentiation among N□O-reducing communities. Moreover, *nirK* was more abundant and the only expressed *nir* gene, which is consistent with recent evidence showing that *nirK* is generally more prevalent than *nirS* across terrestrial ecosystems [59]. This also aligns with site-specific studies indicating that *nirK* denitrifiers rather than *nirS* types, drive N□O production [13, 60], highlighting their distinct ecological roles [59, 61, 62]. This is supported by our metatranscriptomic analyses in which we found no expression of *nirS*-associated helper genes (*nirF, nirN, nirJ, nirE,* and *nirT*), which are essential for a functional catalytic nitrite reductase [63, 64, 65]. *nirS* expression is therefore more energetically demanding and strongly dependent on strict anoxic conditions [60, 66] in contrast to *nirK* that function independently of additional regulatory genes [59, 67, 68], suggesting physiological and habitat preferences and niche partitioning between microbial community harboring nitrite reductases in various environments [61, 69].

Genome-resolved analyses revealed high abundance of partial denitrifiers possessing *nir* but lacking *nosZ*, outcompeting complete denitrifiers (Figure 4E). This is consistent with findings by Pold et al., 2025 [18], who reported that partial denitrifiers initiating but not terminating the denitrification pathway dominate across biomes. Expression of *nirK* and *nosZ* clade I was mostly associated with high abundance of Rhizobiales, which is consistent with denitrifying rhizobia [17]. This pattern reflects the dominance of plant-associated lineages among *nosZ* clade I N□O reducers [58, 70] and potential niche differentiation between *nosZ* clades driven by carbon quality [29]. In particular, members of the family Bradyrhizobiaceae and the order Rhizobiales actively expressed denitrification genes across most surface samples in the present study. These taxa are known to be aerobic or facultatively aerobic denitrifiers [71–75], making them well suited to the oxic–anoxic dynamics in the rhizosphere related to diel patterns in root respiration driven by photosynthesis and the transition zone of surface peat. Our genome-resolved and metatranscriptomic analyses further suggested that N_2_O emissions in this system were predominantly associated with heterotrophic denitrifiers rather than by autotrophic nitrification or nitrifier denitrification pathways. This contrasts earlier reports from Arctic peat soils, where nitrification has been proposed as one of the key control on N□O production by regulating the supply of NO and NO for downstream denitrification, specifically in bare peat during dry years [76, 77]. In the present study, however, genes associated with nitrification were absent from the MAG datasets, and only low-abundance *amoA* genes in ammonia-oxidizing bacteria (AOB) were detected in the metagenomes. Moreover, no transcriptional activity of *amoA* was observed, nor was elevated expression of the downstream nitrification gene hydroxylamine oxidoreductase (*hao*) detected at the transcript level. Together, these results show that nitrifier-driven oxidation processes are unlikely to contribute substantially to N_2_O production in this acidic permafrost peatland, emphasizing the role of heterotrophic denitrification as the primary source of N_2_O and nitrogen gas losses in this ecosystem. This aligns with earlier reports demonstrating that denitrification can produce substantial amounts of N□O in northern peat soils [57, 78]. However, the source of NO supply for denitrification in this system remains speculative. It may be supplied from undetected comammox bacteria or localized internal nitrification in microenvironments not sampled here, such as oxic microsites near root zone, moss layer, or deeper peat layers [78]. We also cannot exclude external sources such as atmospheric deposition through rainfall or hydrological transport likely provide NO in these pristine, unfertilized soils [80, 81].

Availability of labile C is likely one of the drivers behind the diurnal N_2_O variability. With high N_2_O flux along the thaw gradient, daytime CO uptake, nighttime CO and CH release, and all carbon fluxes peaked in the fen, indicates high productivity with enhanced nutrient availability with solute inputs from groundwater and mineral soil [82–85]. This spatial trend in C turnover was well-aligned with decrease in DOC aromaticity. Despite minimal expectations of diel shifts in peat chemistry, DOC, NO , and NH concentrations peaked during the day in fen and palsa moss, matching the strongest diurnal N□O signals, supporting our expectation of higher diurnal variation in habitats with higher content of reactive N and DOC that support the denitrifier communities. The coupling between photosynthetic activity and N_2_O fluxes has been broadly discussed [7] and in our study site likely arises from multiple interacting mechanisms. Labile carbon inputs via root exudation [86], from the actively photosynthesizing moss layer [87], or from photoautotrophic microbes at the mire surface [88] may directly fuel denitrification by providing easily available electron donors. These inputs may also enhance nitrogen mineralization by stimulating extracellular enzyme production [89]. In addition, diel redox fluctuations driven by photosynthetic oxygen release may enhance daytime N mineralization, thereby increasing substrate availability for denitrification [7]. Nevertheless, the relative importance of these processes in driving the diel changes in denitrifier activities remains unresolved and need further investigation.

## Conclusion

While diel N_2_O pattern have been previously reported for terrestrial ecosystems [7, 9], direct molecular evidence for the microbial contribution to this phenomenon has been lacking. By coupling diurnal N□O flux measurements with qPCR, metatranscriptomics, and metagenomics, this study provides mechanistic insight into the microbial drivers of diurnal N□O flux variability in a permafrost thawing gradient in a nutrient-poor peatland. The integrated information from the omics data suggests that elevated daytime N□O emissions were primarily attributed to enhanced microbial N□O production via partial denitrification, whereas lower nighttime fluxes reflected likely resulting from a combination decreased N□O production and enhanced N□O consumption. Despite the inherent biological variability due to the field-based nature of this study, our findings demonstrate that diurnal variation in net N□O fluxes is driven by the balance between microbial N□O production and consumption, primarily through shifts in denitrification gene expression. Microbial activity in the shallow surface layer, often missed by conventional sampling strategies, controlled diel N□O exchange at the surface–atmosphere interface. Extending integrated microbial–flux approaches as in this study across different geographical regions, ecosystem types, seasons, and nutrient gradients will be essential to estimate the role of diel N-cycling microbial dynamics and N_2_O flux and to improve predictions of N□O emissions from rapidly changing northern ecosystems and global N□O budgets.

## Data availability

All raw sequencing data generated and analyzed in this study are publicly available in the NCBI SRA database under BioProject accession PRJNA1481428 (metagenomes) and BioProject accession PRJNA1489742 (metatranscriptomes). The HMMER profiles used in this study and MAG dataset produced in this study are publicly available in Zenodo https://doi.org/10.5281/zenodo.21454293. All MAGs from the EMERGE project are available via Zenodo at https://doi.org/10.5281/zenodo.7596016 [90].

## Acknowledgements

DP acknowledges Erik Lundin and other staff of the Abisko Scientific Research Station for logistic support during sampling. Virginia Rich from the Ohio State University and Ruth Varner from the University of New Hampshire are acknowledged for valuable discussions and sharing site information.

## Funding

The project was financially supported by following funding agencies: the Research Council of Finland in the frame of projects Thaw-N (grant nos. 349503 and 353858), Nitrobiome (grant no. 361980) and N-Perm (grant no. 341348); FWF, Austria, in the frame of PERNO (grant no. M03335) and the Knut and Alice Wallenberg Foundation (grant no. KAW2023.0324). NYT and MG were supported by the European Research Council (ERC) under the European Union’s Horizon 2020 research and innovation programme (grant agreement no. 951288, Q-Arctic).

## Conflicts of interest

The authors declare no conflicts of interest for this manuscript.

## Supplementary files

**Supplementary Figure 1:** Box plot showing daytime and nighttime fluxes of (A) CH_4_ and (B) CO_2_ fluxes across the four habitats. Boxes represent lower and upper quartiles, median (thick black line), smallest and largest values without outliers (thin black line); n=6 independent samples. The dashed horizontal line marks the null expectation (0), where values above and below the line indicate positive and negative deviations, respectively. Statistical differences among habitats were assessed using two-way ANOVA, whereas paired t-tests were used to assess day–night differences within each habitat. Asterisks indicate significant differences between day and night (t-test, ∗*P* <0.05, ∗∗*P* <0.01, ∗∗∗*P* <0.001). Gold and gray colors indicate daytime and nighttime sampling, respectively.

**Supplementary Figure 2:** Soil chemistry at 15 cm depth. Estimation of (A) DOC (dissolved organic carbon); (B) NO_3_ (nitrate); (C) SUVA_254_; (D) NH_4_ (ammonium) from four habitat. Coloured circle points represent means, and vertical whiskers indicate the standard deviation. Statistical differences among habitats were assessed using two-way ANOVA, and between day and night paired *t-*test was applied to assess day-night differences within each habitat (*n*=3). No significant day–night differences were detected (*p* > 0.05). Gold and gray colors indicate daytime and nighttime sampling, respectively.

**Supplementary Figure 3:** Elemental and isotopic composition of carbon (C) and nitrogen (N) measured by EA–IRMS. Estimation of N% is shown for (A) 2 cm and (E) 15 cm depth. Estimation of C% is shown for (B) 2 cm and (F) 15 cm depth. Nitrogen isotope composition (δ^1^ N) is shown for (C) 2 cm and (G) 15 cm depth. Carbon isotope composition (δ^1^³C) is shown for (D) 2 cm and (H) 15 cm depth. Coloured circle points of Dot-and-whisker plot represent means, and vertical whiskers indicate the standard deviation (*n*=3). Statistical differences among habitats were assessed using two-way ANOVA and between day and night paired *t-*test was applied to assess day-night differences within each habitat (*n*=3). No significant day–night differences were detected (*p* > 0.05). Gold and gray colors indicate daytime and nighttime sampling, respectively.

**Supplementary Figure 4:** Estimation of DON (dissolved organic nitrogen) at (A) 2 cm depth and (B) 15 cm depth samples from four habitat. Coloured circle points represent means, and vertical whiskers indicate the standard deviation (*n*=3). Statistical differences among habitats were assessed using two-way ANOVA, and between day and night paired *t-*test was applied to assess day-night differences within each habitat. No significant day–night differences were detected by *t*-tests (*p* > 0.05). Gold and gray colors indicate daytime and nighttime sampling, respectively.

**Supplementary Figure 5:** Correlation analyses between N_2_O flux, nutrient analysis and qPCR-based gene expression at (A) 2 cm depth and (B) 15 cm depth. Asterisks denote statistically significant correlations (∗*P* < 0.05, ∗∗*P* < 0.01; Spearman’s correlation).

**Supplementary Figure 6:** Abundance of bacterial and archaeal 16S rRNA gene across four habitats (Palsa lichen, Palsa moss, Bog and Fen) at two soil depths (2 and 15 cm). Bacterial 16S rRNA gene copy number at DNA level are shown for (A) 2 cm and (B) 15 cm depth. Archaeal 16S rRNA gene copy number at DNA level are shown for (C) 2 cm and (D) 15 cm depth. Bacterial 16S rRNA gene copy number at RNA level are shown for (E) 2 cm and (F) 15 cm depth. Archaeal 16S rRNA gene copy number at RNA level are shown for (G) 2 cm and (H) 15 cm depth. Gene abundances were determined by quantitative PCR (qPCR). Three biological replicates were analyzed for each habitats, with each biological replicate representing the mean of two technical replicates. Data are presented as mean ± standard deviation (SD). Differences in gene abundance between daytime and night-time samples within each habitat were assessed using the *t*-test. No significant day–night differences were detected (*P* > 0.05). Gold and gray colors indicate daytime and nighttime sampling, respectively.

**aaaSupplementary Figure 7**: Abundance of denitrification and N□O-reduction genes across four habitats (Palsa lichen, Palsa moss, Bog and Fen) at two different soil depths (2 and 15 cm). Gene copy number are shown for *nirK* gene at (A) 2 cm and (B) 15 cm, *nirS* gene at (C) 2 cm and (D) 15 cm depth, *nosZ* clade I gene at (E) 2 cm and (F) 15 cm, *nosZ* clade II gene at (G) 2 cm and (H) 15 cm depth. Gene abundances were determined by quantitative PCR (qPCR). Three biological replicates were analyzed for each habitat, with each biological replicate representing the mean of two technical replicates. Data are presented as mean ± standard deviation (SD). Differences in gene abundance between daytime and night-time samples within each habitat were assessed using the *t-* test. No significant day–night differences were detected (*P* >0.05). Gold and gray indicate daytime and nighttime samples, respectively.

**Supplementary Figure 8:** Expression of N□O reductase and NO-forming nitrite reductase genes in the four habitats (Palsa lichen, Palsa moss, Bog and Fen) based on qPCR analysis of soils at 15 cm depth. (A) Habitat-specific and (B) overall *nirK*/*nosZ* expression ratios, representing the balance between the potential for N□O production and N□O reduction. (C) Expression of N□O reductase gene *nosZ* clade I and (D) expression of nitrite reductase gene *nirK*. Three biological replicates were analyzed for each habitat, with each biological replicate representing the mean of two technical replicates. Data are presented as mean ± standard deviation (SD). Differences in gene expression and *nirK/nosZ* expression ratios between daytime and nighttime samples, both within individual habitats and across all habitats were assessed using *t*-tests. No significant day–night differences were detected (*P* >0.05). Gold and gray indicate daytime and nighttime samples, respectively.

**Supplementary Table 1:** Details of qPCR primers and thermal cycling conditions used in this study.

**Supplementary Table 2:** N□O fluxes and ancillary environmental variables measurement during flux measurements.

**Supplementary Table 3:** Soil chemical characteristics of the sub-arctic permafrost habitats during day-night sampling

**Supplementary Table 4:** Taxonomic distribution of *nirK* and *nosZ* gene distribution at different taxonomic levels based on metagenome analysis from both soil depths

**Supplementary Table 5:** Expression of nitrogen cycling genes in daytime and night-time samples based on metatranscriptomic analyses of the arctic permafrost habitats.

**Supplementary Table 6:** Distribution of nitrogen cycling genes within metagenome assembled genomes (MAGs)

**Supplementary Table 7:** Abundance of nitrogen cycling genes inferred from metagenomic sequence analysis.

**Supplementary Table 8:** Taxonomic distribution of complete and partial dentification containing genes at MAGs (metagenome assembled genome) level

## Notes

### Competing Interest Statement

The authors have declared no competing interest.

### Summary of Updates

This version of the manuscript update with the metatranscriptomics analysis to provide more depth interpretation with updates figures

