## Supplementary figures and images for "Denitrifier activity drives diurnal N_2_O flux dynamics in a thawing arctic permafrost"

### Fig_S1.tiff

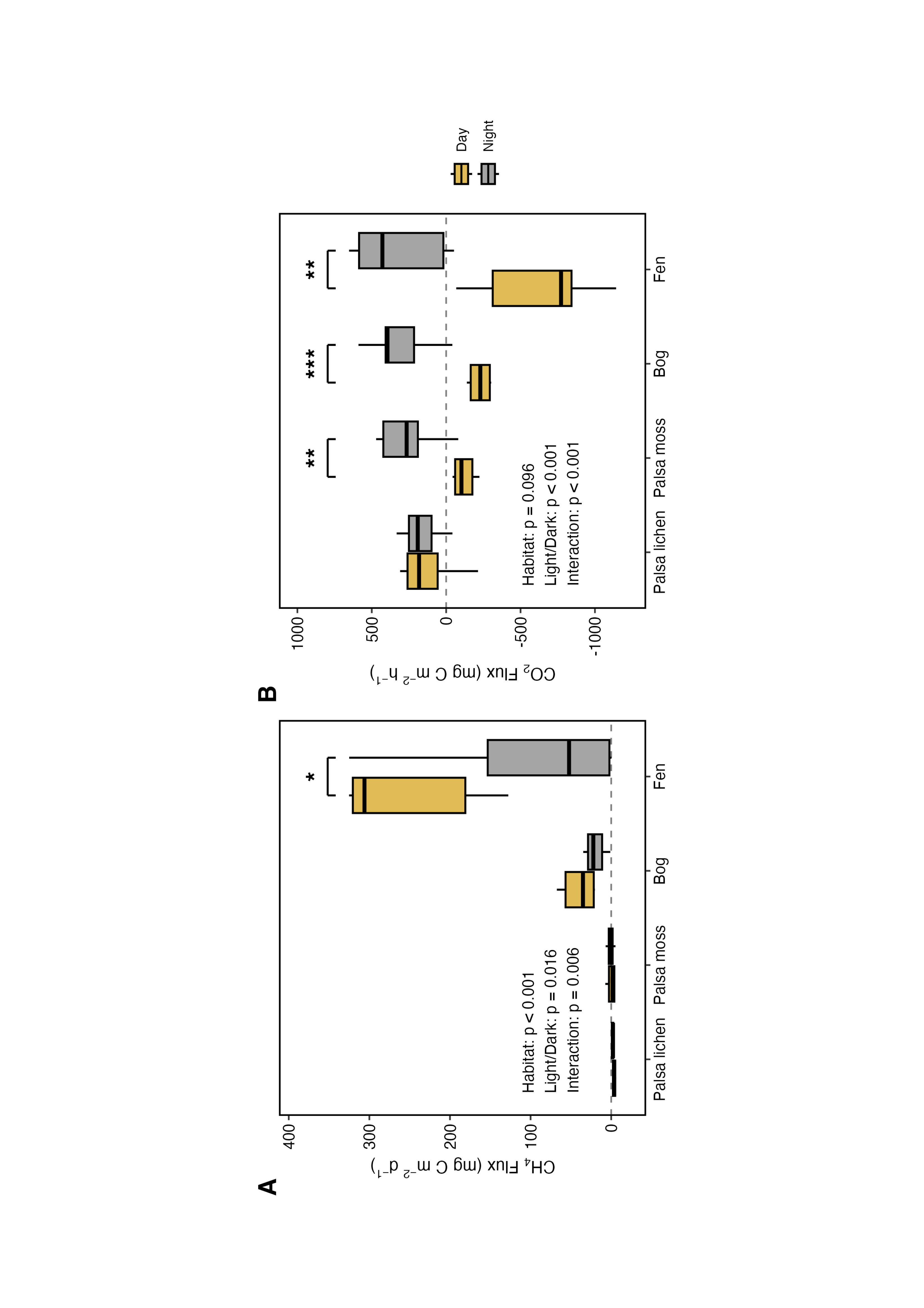

### Fig_S2.tiff

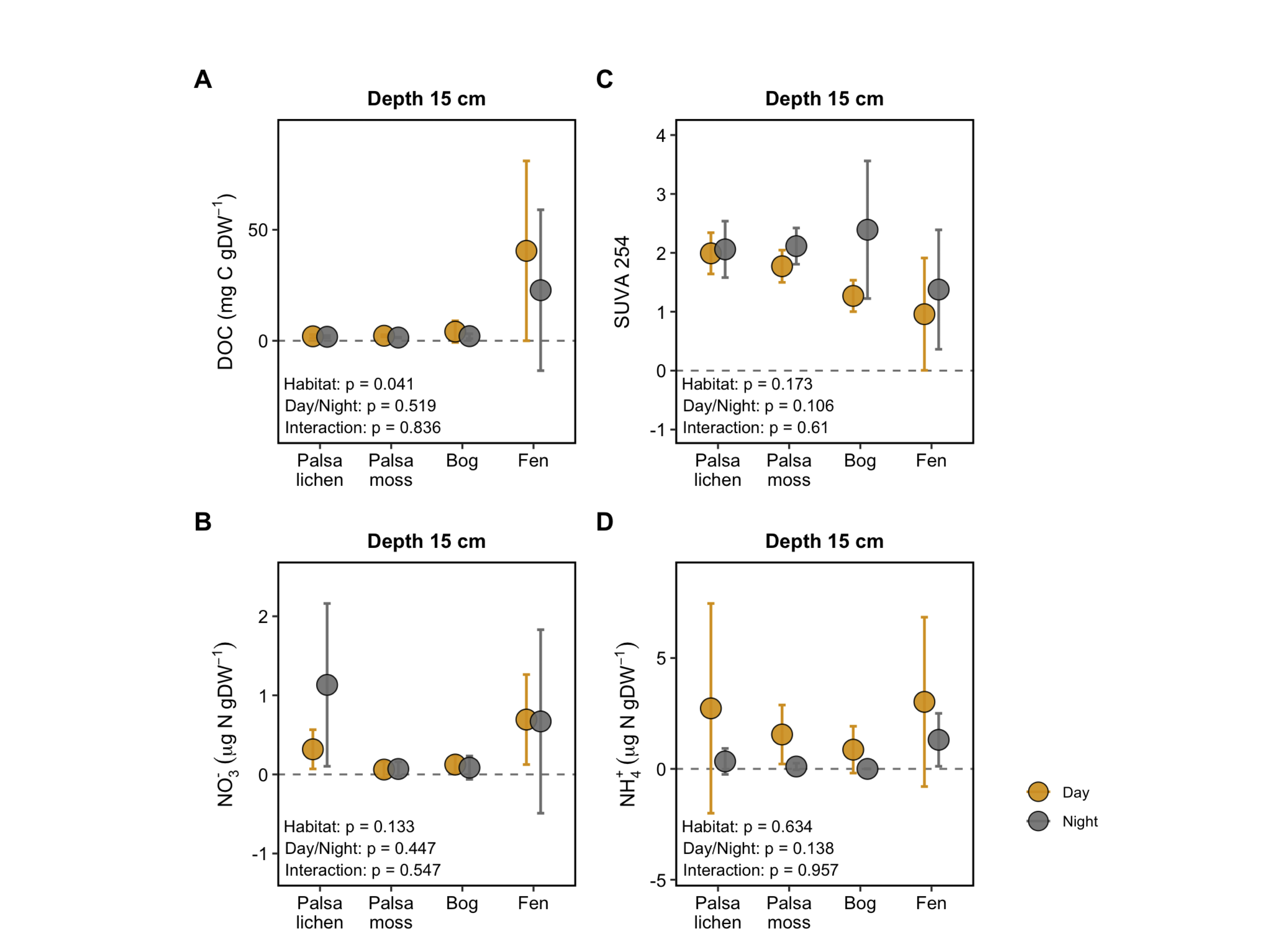

### Fig_S3.tiff

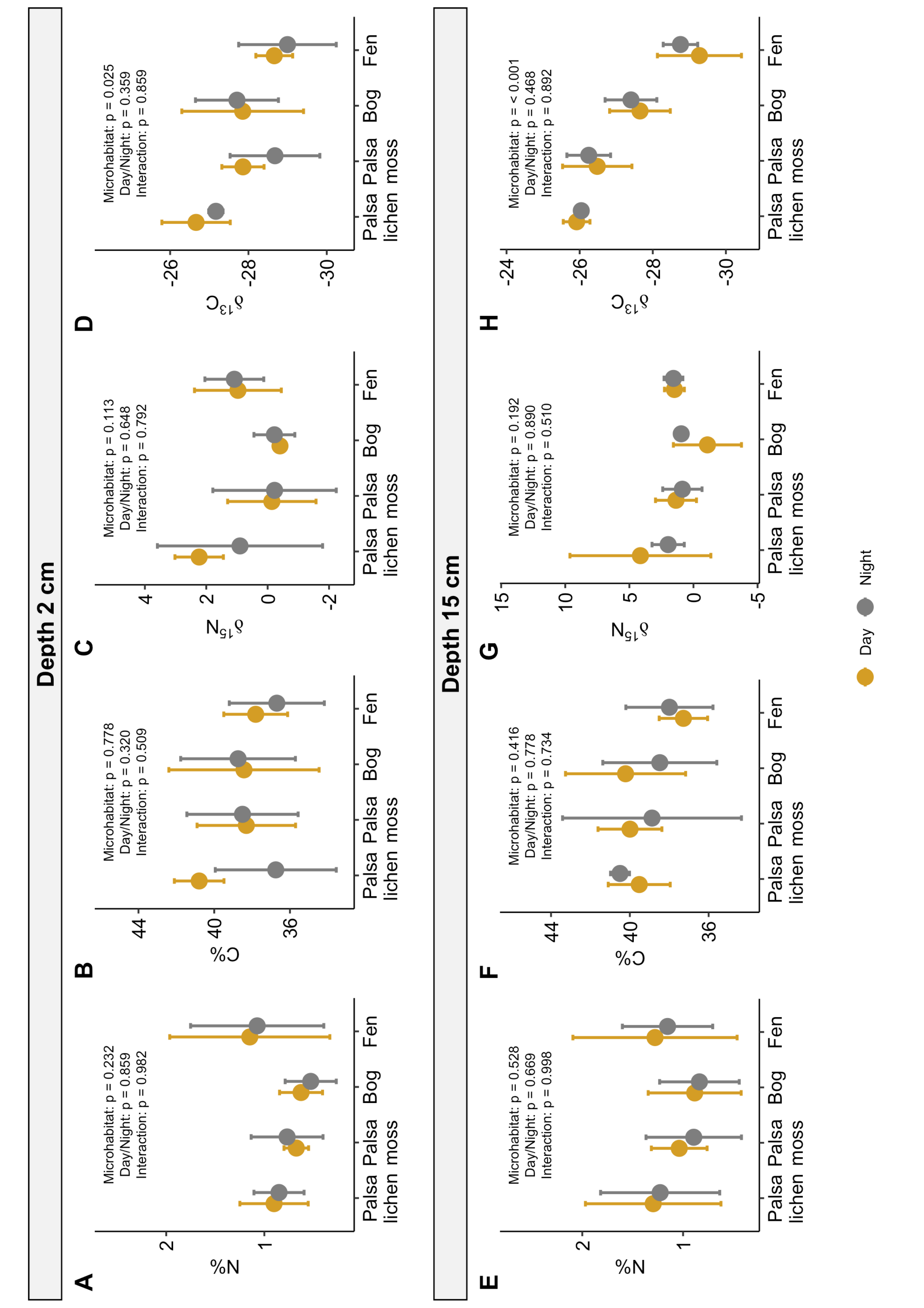

### Fig_S4.tiff

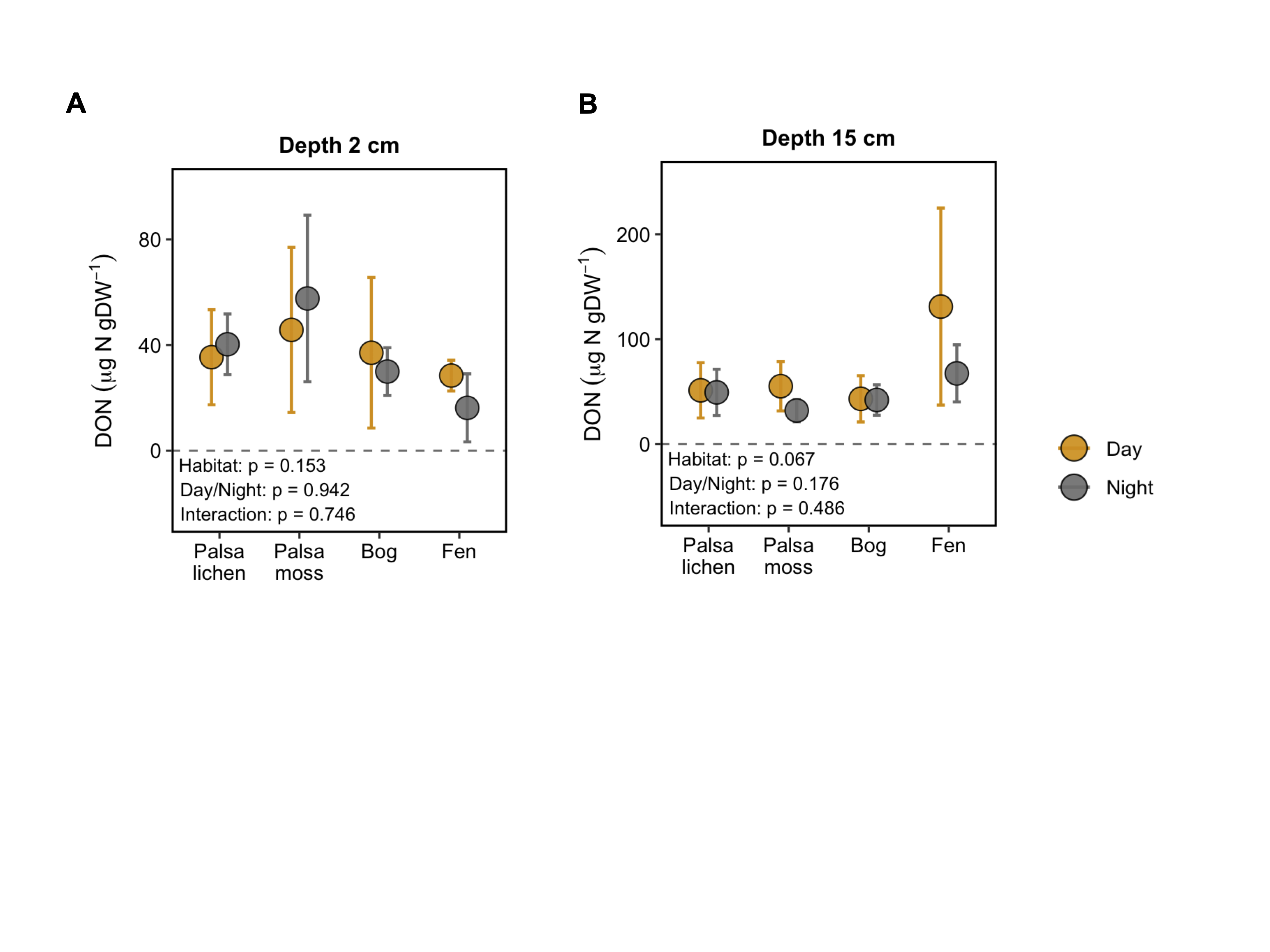

### Fig_S5.tiff

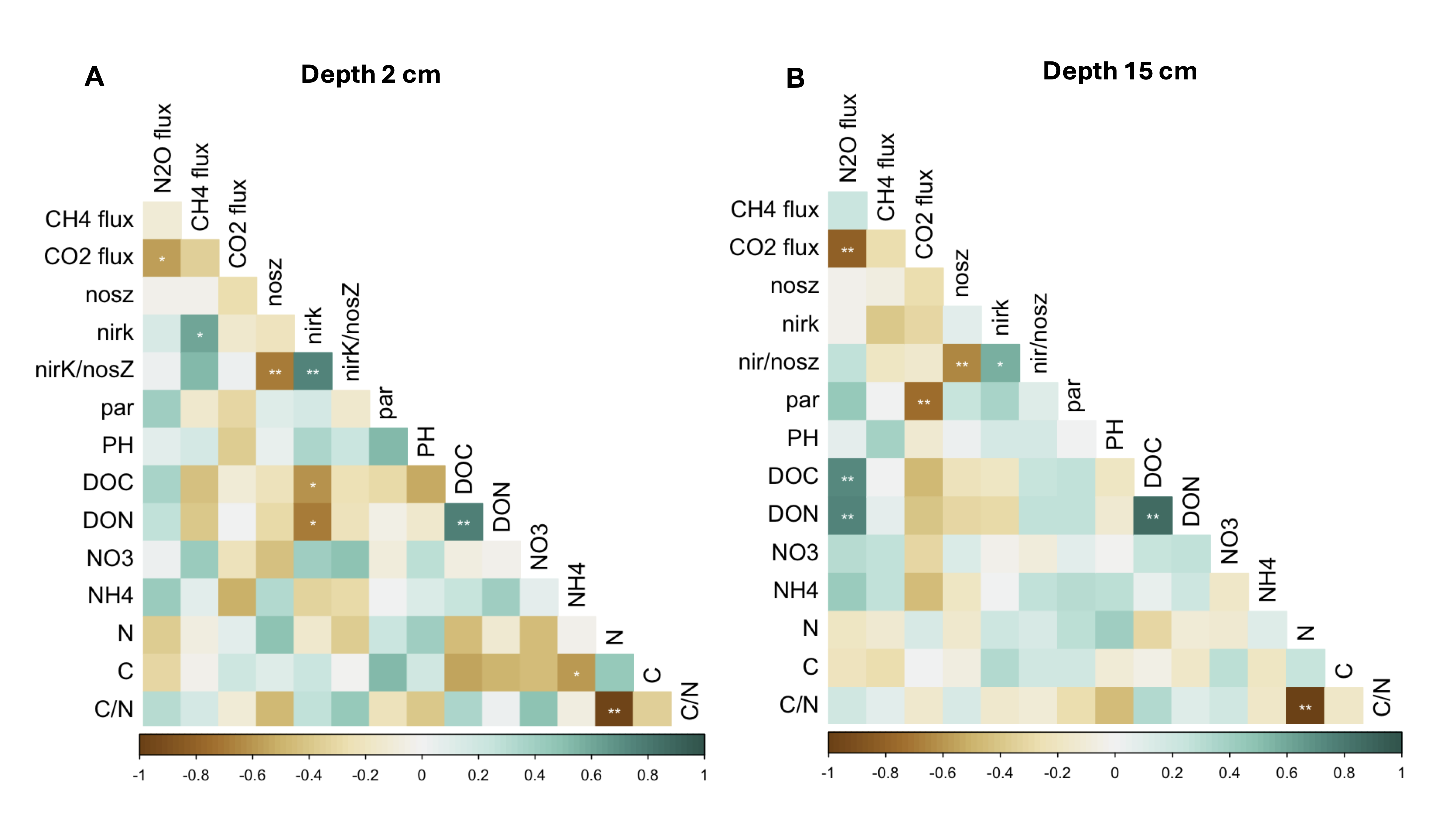

### Fig_S6.tiff

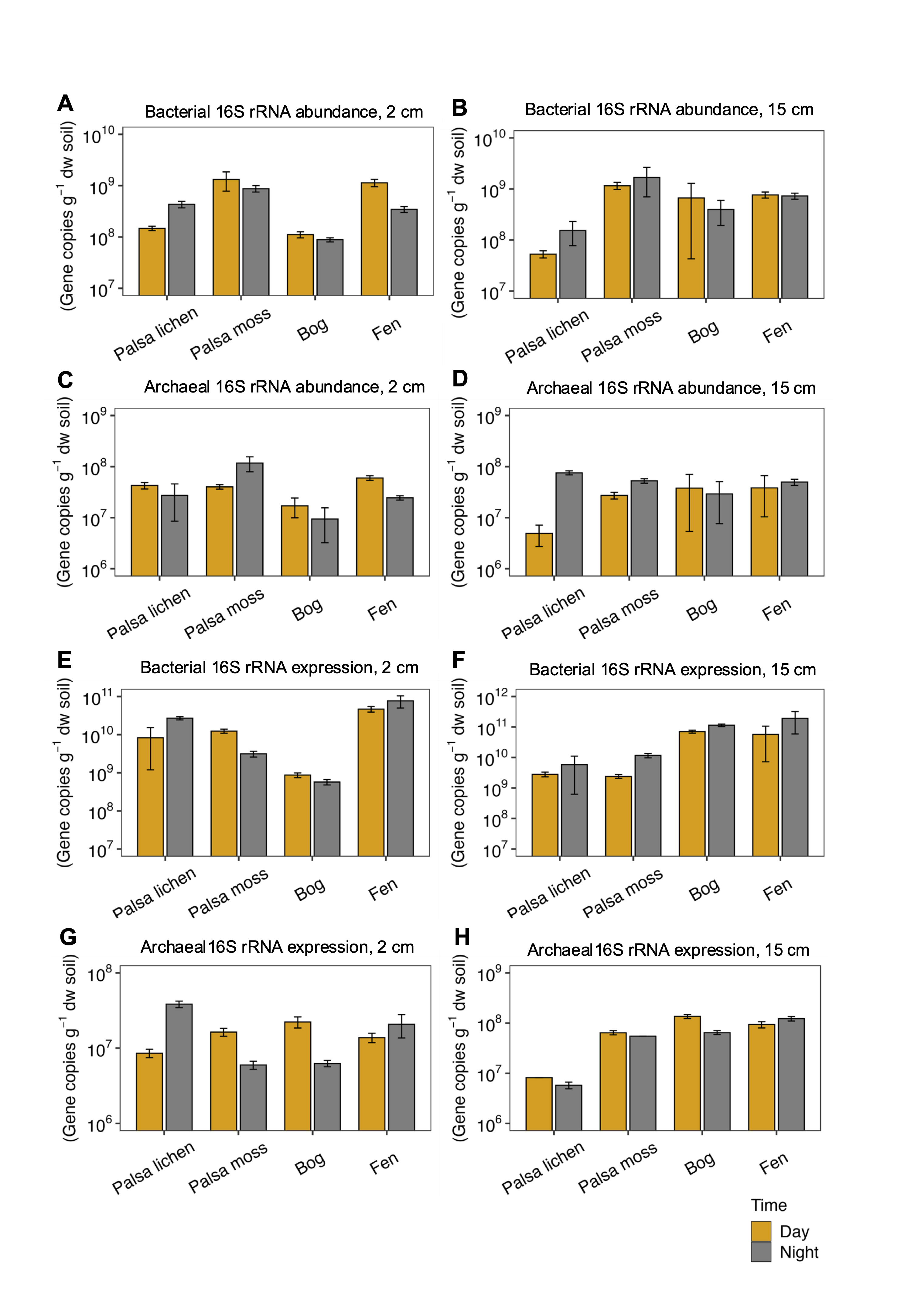

### Fig_S7.tiff

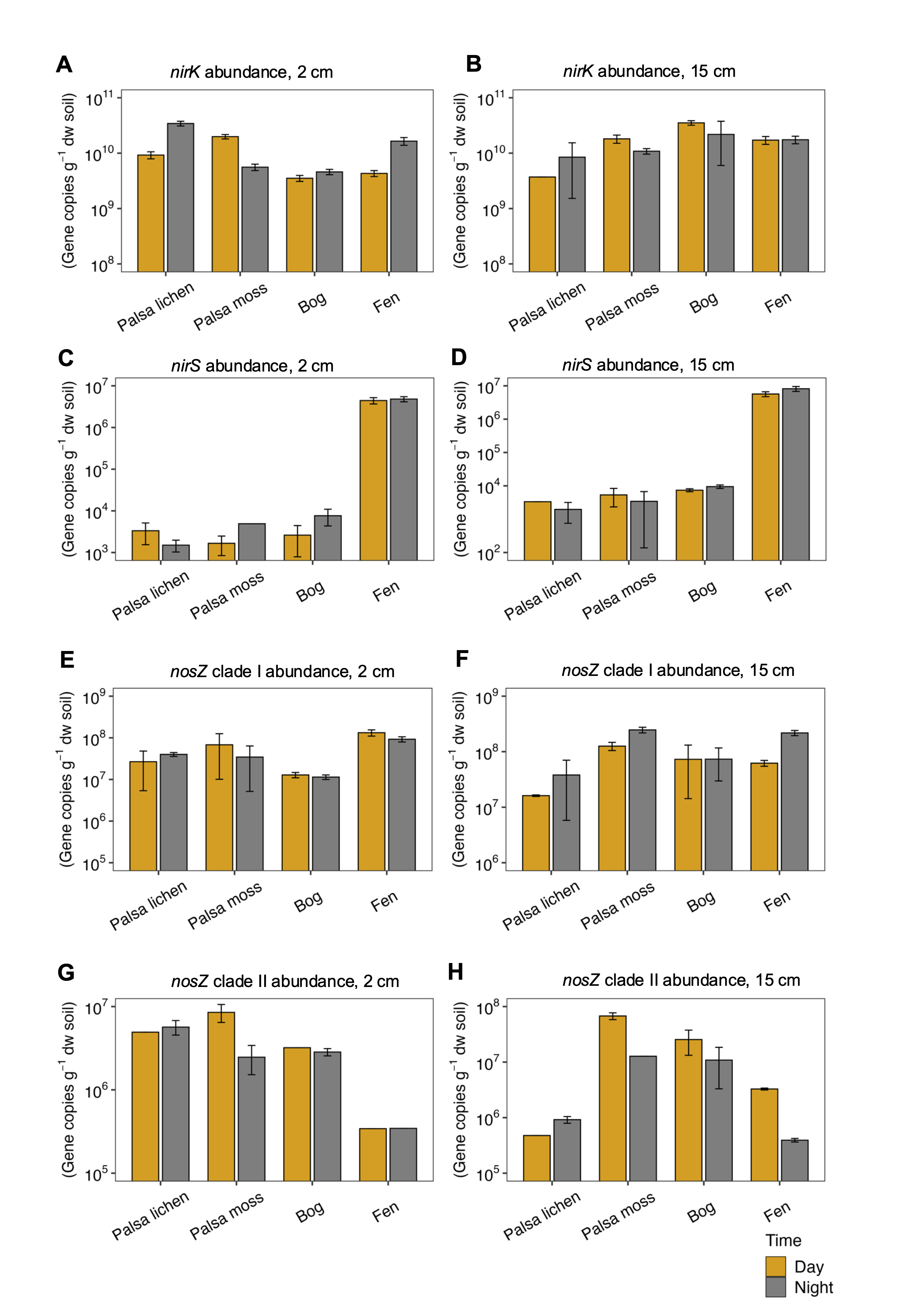

### Fig_S8.tiff

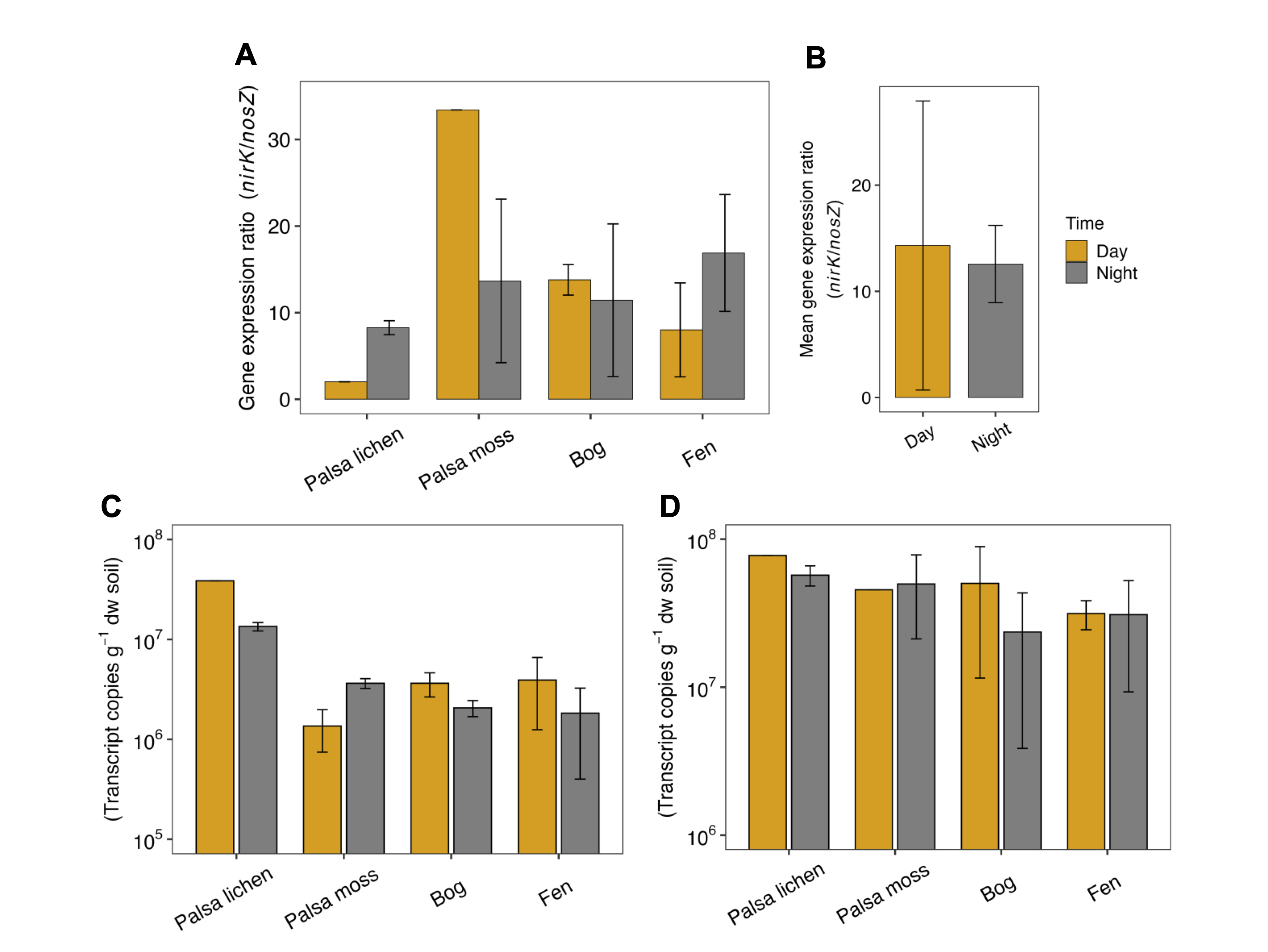
