## Supplementary material for "Denitrifier activity drives diurnal N_2_O flux dynamics in a thawing arctic permafrost": supplymentary methods

**Sampling site description**

Our study Stordalen Mire (68°35’N, 19°04’E), a ~25 hectares permafrost peatland located 10 kilometers east of the Abisko Natural Sciences Research Station. It represents an arctic eastern continental peatlands system, developed on permafrost, with its central area covering around 15 hectares [1]. The region is in the southern border of permafrost occurrence, and isolated permafrost is currently present in the dry uplifted areas on the peatland (palsa), but it is threatened by ongoing permafrost thaw [2].

The Stordalen mire ecosystem has three distinct habitats with variable permafrost and moisture conditions, representing a gradient of thaw intensity. The first is the palsa, ombrotrophic peat surfaces uplifted by a permafrost core. The vegetation is primarily dominated by lichens (Cladonia spp.) and shrubs, such as *Empetrum hermaphroditum*, *Betula nana*, *Vaccinium uliginosum*, *Vaccinium vitis-idaea*, and *Rubus chamaemorus*. Additionally, mosses like *Dicranum elongatum* and *Sphagnum fuscum* are also present. In the current study, we divided the palsa further two categories: moss-dominated areas, wetter area (palsa moss) and lichen-dominated drier areas (palsa lichen). The second habitat in the thaw gradient is the bog, characterized by a series of hollows within the palsa, which are filled with stagnant, ombrotrophic water. This environment supports a rich diversity of graminoid species, including *Sphagnum balticum, Sphagnum lindbergii*, and *Sphagnum riparium*, as well as vascular plant sedges such as *Carex rotundata* and *Carex rostrata*. The thirds habitat is the fen, situated in the lower regions of the mire, where free-flowing water promotes the growth of large aquatic graminoid species, including *Sphagnum balticum*, *Sphagnum lindbergii*, and *Sphagnum riparium,* along with vascular plants like cotton grass (*Eriophorum vaginatum* and *Eriophorum angustifolium*).

**N_2_O flux measurment**

To measure N_2_O flux, we paired a transparent acryl-glass chamber (height 25 cm, diameter 25 cm) with a state-of-the-art portable gas analyzer (Aeris MIRA Ultra N_2_O/CO_2_) with a precision of less than 2 ppb for N_2_O. For the first night measurements (12 flux measurement) another portable laser Li-7820 (Licor) was used due to unavailability of the default PGA. Preinstalled chamber base collars (PVC) with a dimeter of 25 cm were used. Duplicate measurements were subsequently conducted at each plot for a robust flux estimate. We measured gas concentrations in the chamber headspace at 1-second intervals over a 5-minute enclosure period. Auxiliary data including, chamber temperature, air pressure, relative humidity and PAR were recorded during flux measurements as previously described [3]. Hand-held meters were used for soil temperature at 2 and 15 cm and soil moisture as volumetric water content at 0-6 cm (HH2 with ML2x ThetaProbe, Delta-T Devices). For N_2_O flux calculation we followed our previously published work [3].

For N_2_O flux calculation we followed our previously published work [3]. Briefly, we used the goFlux package in R (v 0.2.0., Rheault 2025) [4], which uses both linear models (LM) and nonlinear (HM) and gives suggestion of recommended estimation method for each flux measurement. This open-source tool is specifically designed to work with high throughput, laser-based GHG data. We calculated fluxes by utilizing all data points from the 5-min chamber closure, expect for the first 8 seconds, which were excluded. The measured data on temperature, pressure, and relative humidity were used for the calculation.

**Soil geochemical analysis**

We determined the content of water extractable ammonium (NH_4_^+^), various anions, including nitrite (NO_2_^‒^), nitrate (NO_3_^‒^), sulfate (SO_4_^²‒^), phosphate (PO_4_³^‒^), and chloride (Cl^‒^), and dissolved organic carbon (DOC) and dissolved nitrogen (DN) content based on water extraction using a soil-to-water ratio of 3:10 (v/v). Extracts were prefiltered through filter paper (Whatman, no. 42) and final filtering was done with 0.45 µm syringe filters. Anion concentrations in water extracts were analyzed using ion chromatography (Dionex ICS-2100, Thermo Scientific), DOC and DN, on a TOC-V CPH analyzer (Shimadzu Scientific). The dissolved organic N (DON) content was derived by subtracting water soluble mineral N species (NO_2_^‒^, NO_3_ ^‒^, and NH_4_^+^) from DN. For all analyses, extraction blanks (n = 3) were subtracted from the sample concentrations. For total C and N content and C:N ratio, and δ^13^C and δ^15^N, oven-dried samples were ground by a ball mill and analyzed on an EA-IRMS (Elemental Analyzer Isotope Ratio Mass Spectrometer; Delta XP Plus 191 IRMS with Flash EA 1112 Series Elemental Analyzer and a Conflow III open split interface, Thermo Finnigan). Specific UV absorbance at 254 nm (SUVA_254_) was measured with a laboratory benchtop spectrophotometer (UV-1800, Shimadzu) and used for estimation of DOC aromaticity [5].

**Metatranscriptomic data analysis**

The quality of the raw paired-end reads was assessed using FastQC (Andrews, 2010). Sequences of good quality (Q > 30) were selected for further analysis using Trimmomatic [6]. Ribosomal RNA sequences were removed using SortMeRNA (v4.3.6) [7] against the SILVA rRNA database (release 138) with default parameters. In total, approximately 1.35 billion high-quality non-rRNA reads were obtained for downstream analyses. The resulting high-quality non-rRNA reads were assembled de novo using rnaSPAdes (v3.15.5) [8]. Open reading frames (ORFs) were predicted from assembled contigs using Prodigal (v2.6.3) [9] in metagenomic mode (-p meta), and the resulting coding sequences were functionally annotated using eggNOG-mapper (v2.1.12) against the KEGG, COG and pfm database with an E-value cutoff of 1e−5. Filtered reads from each sample were mapped to the predicted coding sequence library using Bowtie2 (v2.5.1) [10] with default parameters. Resulting SAM files were converted, sorted, and indexed using SAMtools (v1.17) [11]. Gene-level read counts were obtained from the alignments, and expression levels were normalized as transcripts per million (TPM), accounting for gene length and sequencing depth. In addition, HMMER profiles for the *nir* and *nosZ* genes were used to identify corresponding active funtional gene sequences from quality-filtered reads using a maximum E-value threshold of 1 × 10⁻⁵. The retrieved sequences were subsequently analysed with GraftM [12] using reference phylogenies for *nosZ* [13] and *nir* [14] to determine the taxonomic affiliation and relative abundance of *nosZ* and *nir* genes.

**Metagenome assembly and analysis**

Quality‑filtered reads were assembled using the MEGAHIT assembler v1.2.9 [15] with parameters (--k-min 21 –k-max 141 –k-step 12). Metagenome‑assembled genomes (MAGs) were reconstructed by applying three binning algorithms: MaxBin2 v2.2.7 [16], MetaBAT2 v2.15 [17], and CONCOCT v1.1.0 [18]. Resulting bins were consolidated using DASTool v1.1.5 [19] and depelicated using  dRep [20]. Quality of the MAG was checked yúsing CheckM [21]. Taxonomic classification of MAGs was performed using the *classify_wf* workflow of GTDB‑Tk (v2.4.0) with the GTDB R220 release [22].

ORFs were predicted from metagenome assemblies using Prodigal (v2.6.3, meta mode) and functionally annotated with eggNOG‑mapper (v2.1.12; eggNOG v5.0; E‑value ≤1e−5) against the KEGG, COG and pfm database. Filtered reads were mapped to predicted genes using Bowtie2, processed with SAMtools, and normalized to count per million. Taxonomic affiliations and relative abundances of *nir* [14] and *nosZ* [13] genes from metagenome were inferred using GraftM [12] and with established *nos*Z and *nir*K reference phylogenies.

<https://doi.org/10.1093/bioinformatics/btu170>
